# Scalable Batch Fabrication of Ultrathin Flexible Neural Probes using Bioresorbable Silk Layer

**DOI:** 10.1101/2021.04.19.440418

**Authors:** Clement Cointe, Adrian Laborde, Lionel G Nowak, David Bourrier, Christian Bergaud, Ali Maziz

## Abstract

Flexible deep brain probes have been the focus of many research works and aims at achieving better compliance with the surrounding brain tissue while maintaining minimal rejection. Strategies have been explored to find the best way to implant a flexible probe in the brain, while maintaining its flexibility once positioned in the cortex. Here, we present a novel and versatile scalable batch fabrication approach to deliver ultra-thin and flexible penetrating neural probe consisting of a silk-parylene bilayer. The biodegradable silk layer provides a temporary and programmable stiffener to ensure ease of insertion of the ultrathin parylene-based flexible devices. The innovative and yet robust batch fabrication technology allows complete design freedom of the neural probe in terms of materials, size, shape and thickness. These results provide a novel technological solution for implanting ultra-flexible and ultrathin devices, which possesses great potential for brain research.

## Introduction

Chronically implanted microelectrodes have been a key tool in neuroscience research by allowing the recording of electrical brain activity at the level of small population of neurons (local field potential, multiunit spiking activity) and of individual neurons (single-unit activity). The past decades have seen impressive technological developments of neural implants incorporating electrodes at the micrometer scale *e*.*g*. silicon-based pin (Utah), flat (Michigan) or wire (floating microwire arrays) for characterization of neuronal activity^1-3^ Such devices are now routinely used in animal studies^4^. Although long-lasting recording was sometimes achieved using these probes ^5^, large variations in electrical recording capability were often reported^6^. The implementation of long-lasting intra-cerebral recordings is limited by the lack of stability at the interface between the conventional electrodes and the brain tissue^7-8^. This is partly due to the mechanical mismatch between the stiffness of the materials, *e*.*g*. silicon, glass, platinum or iridium (E ≈ 150 GPa), constituting such probes and the softness of the cerebral tissue (E ≈ 10 kPa).^9^ This mechanical mismatch which can be as large as seven orders of magnitude leads to irreversible tissue damage and glia scar formation resulting in failure of the device within months or even weeks after implantation^10-11^.

To improve the brain tissue-electrode interface, the use of flexible depth probes has been the focus of many research works and aims at achieving better compliance with the surrounding neural tissue while maintaining minimal rejection^10, 12^. The fabrication of these compliant devices typically involves either the use of soft polymeric materials as substrate *e*.*g*. parylene^13-15^, polyimide^16-17^, polydimethylsiloxane (PDMS)^18^, hydrogels^19^ and/or the use of significantly thinner stiff materials ^20-21^. However, an important issue with flexible neural probes is that they tend to fail penetrating the pia mater and reach their location goal. Indeed, a device that is too soft tends to bend when pressed against a rigid surface, such as the brain pia matter.^12^ Strategies have been explored to find the best way to implant a flexible probe in the brain, while maintaining its flexibility once positioned in the brain. Some teams focus on the use of a stiff shuttle^20, 22-23^ that is removed immediately after implantation, while others promote the integration of a stiff bioresorbable coating that is not removed but dissolves inside the brain in a time scale of minutes to days^24-25^. Due to the tissue trauma caused by implanting and withdrawing the stiff shuttle, the integration of bioresorbable coatings as a temporary stiffener has been shown to better address both mechanical and biological failures^26^. Various bioresorbable polymers *e*.*g*. PEG, PLA, Chitosan and Silk fibroin have been reported to be excellent candidates to add to the polymeric implants to facilitate insertion into the brain^12^. Besides, they benefit to some extent from common attributes, such as high Young’s modulus, proven biocompatibility for *in vivo* purposes, and bioresorption when in contact with physiological tissues^27^.

Even though the reported flexible implants incorporating a biodegradable coating showed successful short-term electrical recordings, the insertion of the coated probe produces an initial surgical trauma in the range of hundreds of micrometers to millimeters^24-25^, a spot size obviously much larger in diameter than that of the probe itself. The biodegradable stiffener coating, nowadays used for stiffening the flexible probes, is incompatible with standard microfabrication techniques and requires complex manual handling procedures that limit further downscaling (less than 10 μm thick) and make it difficult and time-consuming to create many devices in parallel^12, 28-29^. Furthermore, existing flexible probes require additional preparation of carrier support for the biodegradable coating, with a manual assembly procedure which increases the difficulty of reducing the dimensions of the probe and scaling-up the fabrication. A global rethinking of the production framework is therefore needed to effectively achieve a facile integration of biodegradable coatings and allow further development of minimally invasive neural probes.

To achieve this goal, we report here a versatile fabrication framework utilizing a bioresorbable silk fibroin layer that can be integrated in a microfabrication process for preparing ultrathin parylene-based penetrating probes. The probes consist of a silk-parylene bilayer, which was obtained by successively depositing layers on the top of each other. The first layer was obtained using degradable silk fibroin coating as a temporary stiffener that can be used to insert ultrathin parylene-based flexible devices deep in brain. The additional subtlety of the process derives from the exposure of the silk fibroin layer to methanol that increases the crystallized domains in the film allowing the programmable degradation of the stiffening layer through proteolytic reactions. Furthermore, *in vitro* insertion trials in brain phantom showed neither buckling issues of the probe nor undesired alteration of the electrical properties. Finally, high-quality signal recordings *in vivo* in mouse brain slices is demonstrated.

## Results and Discussion

### Development of flexible penetrating neural probe

Our ultrathin flexible probes were developed using standard microsystems techniques. The probes were made of a silk-parylene bilayer that comprises pairs of gold microelectrodes coated with the conducting polymer poly(3,4-ethylenedioxythio-phene):poly(styrene-sulfonate) (PEDOT:PSS) to lower the impedance and obtain a better signal-to-noise ratio for neuron’s recording^13^. The schematics of the fabrication procedures and the probe structure are shown in **Fig. 1A**. Briefly, there are three main fabrication stages. In stage I, a cellulose acetate (CA)-coated glass substrate is first coated with a biodegradable silk fibroin protein before (Stage II) an ultrathin-film parylene-based device structure is microfabricated on top of the silk layer using standard top-down lithography techniques (discussed in more details later in the paper). In final stage III, the whole silk-parylene probe is micro shaped by a Reactive Ion Etching (RIE) process, and released from the glass substrate.

**Figure 1.**
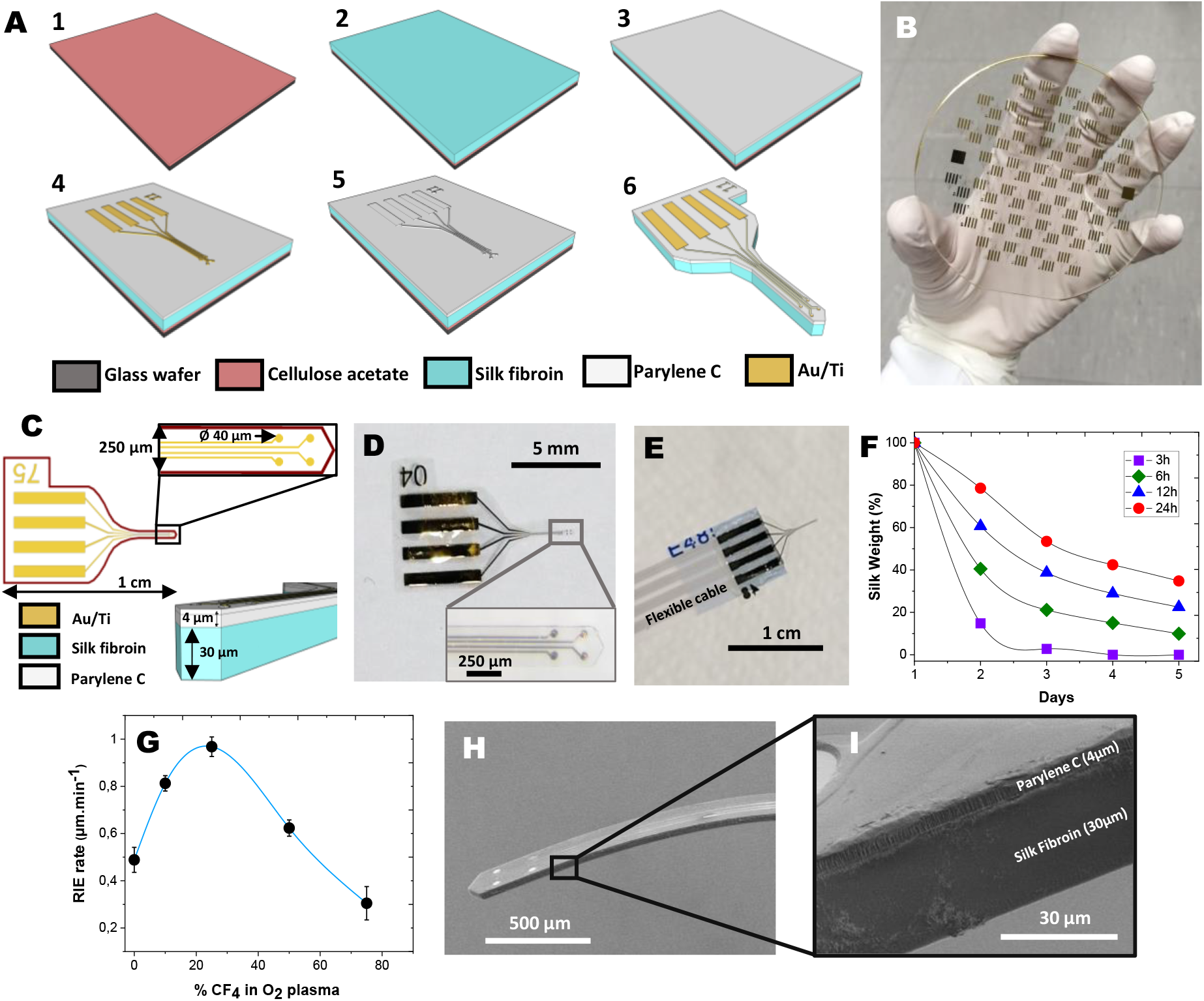
Fabrication of the bilayered silk-parylene neural probes. **A**) Schematic illustration of the main fabrication steps on glass substrate. **B**) Picture of the 4 inches glass substrate after batch microfabrication. The substrate contains 80 elements (less than 1 cm each). **C**) Schematic illustration of the bilayered silk-parylene probe with characteristic dimensions. The device contains four recording microelectrodes of 40 μm in diameter, patterned on a short 3 mm-long and 250 μm-wide shank. **D**) Picture of the microfabricated implant, the inset shows the corresponding zoom-in on the 4 gold microelectrodes. **E**) Fabricated silk-parylene neural probe highlighting the contact pads bonded with FFC cable for the following electrochemical and electrophysiological measurements. **F**) Enzymatic degradation of silk fibroin for different immersion times in methanol, inducing the protein crystallization. **G**) Etching rate at 500 W and 20 mTorr of silk fibroin *vs*. percent of CF4 in O2 plasma. An optimal ratio of 25/75 was found. **H**) SEM picture of the final bilayered silk-parylene shank, and **I**) the corresponding zoom-in SEM image.

There is a couple of key considerations in this design. First, the silk fibroin material is chosen among other bioresorbable materials on the basis of its excellent biocompatibility, tunable biodegradability and high mechanical strength^30^. Indeed, silk fibroin possesses a Young’s modulus E of ∼ 3 GPa so that the critical thickness of the parylene-C probe can be drastically reduced to the µm level, while keeping the handling and the implantation of the probe possible. The biodegradable silk layer is described in detail below because of the specific chemical structure and its importance in the final device. Active research is currently underway on the use of silk biomaterial, in particular to obtain a biodegradable coating for impalpable neural devices^12^. However, no work has yet been reported on the integration of the degradable biopolymer in the batch fabrication of neural devices allowing further scaling down (less than 10 μm thick) and preventing *de facto* any manual handling procedure and buckling issues during implantation.

The thickness of the silk fibroin can be tuned if needed, by adjusting the polymer dilution percentage or the volume of the solution on the casting area accordingly. As an example, a dose of 0.1 mL/cm^2^ of 7 wt. % silk fibroin solution produces films ∼30 ± 5 μm thick. Depending on the surgical procedure, the probe position might need to be adjusted for a prescribed time of several minutes or more, during which it must remain stiff. This critical time is linked to the material degradation rate of the bioresorbable coating. The lifetime of the bioresorbable layer was specifically adjusted, chosen via the crystallinity of the silk protein *i*.*e*. beta-sheet content^31^. Indeed, treating silk by immersion in methanol increases the β-sheet content in the film that will allow for a water-stable structure. In a proteolytic medium, silk fibroin samples degrade at different rates depending on the methanol time treatment: films treated for 3 h and 6 h degrade within a few hours and after up to one week for 12 h and 24 h treatment (**Fig. 1F**). This control of degradation rate is consistent with the literature, where reports have shown proteolytic degradation *in vitro* of water-stable silk films after approximately two weeks^25, 31^.

Next, an ultra-flexible parylene-based neural probe is fabricated on the top of the silk fibroin substrate (**Fig. 1A**) (more details are available in Materials and methods). The parylene is chosen as the substrate for its well-documented biocompatibility, chemical inertness, and high electrical and moisture insulation properties. **Fig. 1 C** shows the design of the ultra-flexible parylene-based neural probe consisting of three layers: the top parylene layer (thickness: 1 μm) for encapsulation, the middle Au layer (thickness: 200 nm) for electrophysiological measurement, and the bottom parylene layer (thickness: 3 μm) for mechanical support. The total thickness of the probe is 4 μm only.

Dry etching was then performed through RIE using a 75/25 O_2_/ CF_4_ gas ratio. In these conditions, as illustrated in **Fig. 1 D, H and I**, microfabrication of bilayered silk-parylene microstructures can be performed with precise control of the geometry, size and shape. The proof of concept presented in **Fig. 1B** contains 80 elements, each device contains four recording gold microelectrodes of 40 μm in diameter, patterned on a short 3 mm-long and 250 μm-wide probes. In other words, large-scale batch fabrication of precisely defined ultrathin silk-parylene probes can be performed directly on the glass substrate. Finally, the dissolution of the cellulose acetate sacrificial layer in acetone was achieved to release the bilayered silk-parylene probes from the substrate (**Fig. 1D**).

### Insertion testing in brain phantom

Our bilayered neural probes are designed to accurately implant ultrathin flexible devices while avoiding buckling during insertion. In addition, the probe was robust enough to easily bond with external electrical connections directly (**Fig. 1E**). Another important parameter lies in the mechanical stability of the stiffening resorbable coating during implantation/explantation process. To illustrate the feasibility of the proposed insertion strategy, cycled insertions of the ultrathin probe into a brain-tissue-mimicking phantom (agarose gel 1 wt. %, Young’s modulus 40 kPa) were performed. The bilayered silk-parylene probes succeeded in penetrating the brain phantom without any buckling or bending issue during the insertion process, which clearly indicates that the proposed insertion shuttle is stiff enough to penetrate the brain phantom. **Fig. 2B** shows the whole process of implanting the thin probe into the brain-tissue-mimicking gel, showing (*1*) the thin probe mounted on the manipulator approaching and (*2*) contacting the gel, (*3*) the probe successfully perforating the gel, (*4*) the thin probe inserted in the gel, (*5* and *6*) the thin probe is deeply inserted into the tissue-mimicking gel. From the insertion experiment (**Fig. 2A**), the first peak force (stage 3) was 0.7 mN indicating the minimum force required to penetrate the tissue-mimicking gel. Other works also showed similar behaviors for the requested minimum force^25, 32^. As **Movie S1** in the Supporting Information shows, the thin probe is explanted from the gel without mechanical damage and successfully reinserted in the gel a second time. These tests give a clear evidence of the reliability of the developed devices for practical surgical handling. It is worth mentioning that the bare parylene shank itself, being only 4 µm thick, curled and could not be handled or manipulated without the bioresorbable silk polymer support.

**Figure 2.**
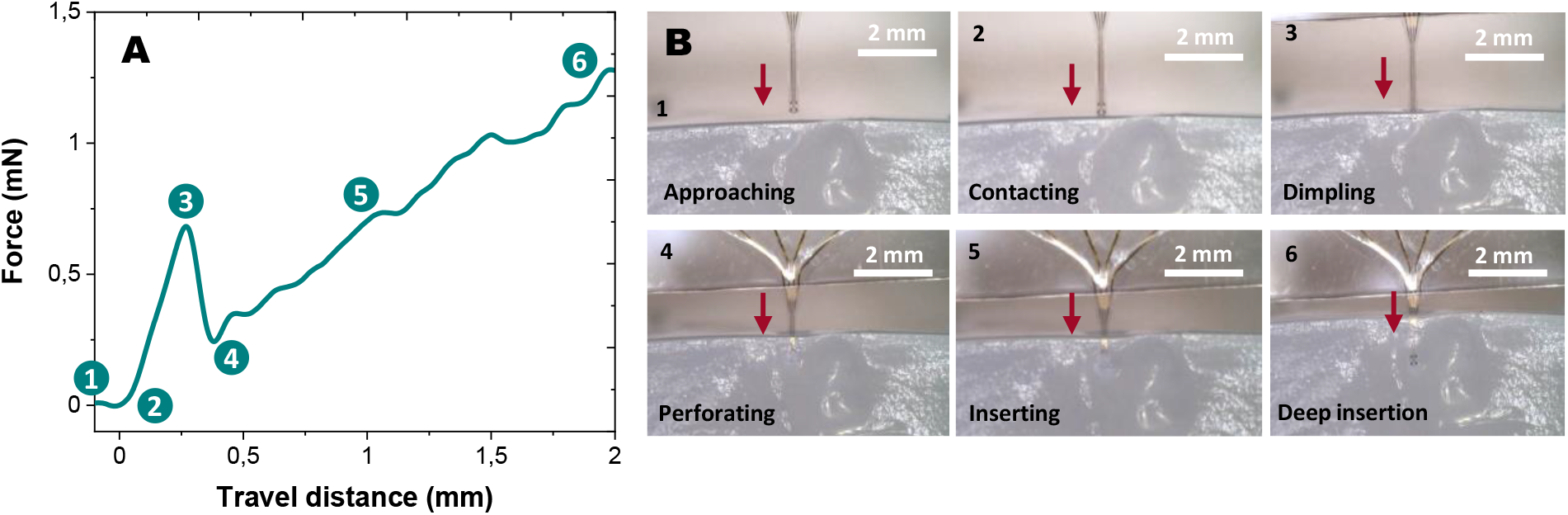
Insertion test of the silk-parylene flexible probe in brain phantoms. **A**) Force profile during insertion of the shank in agarose gel 1 wt. %, Young’s modulus 40 kPa, numbers in correspondence with (B). **B**) Optical pictures showing the different stages of insertion in correlation with the evolution of the force.

### Standard compression tests

We further studied the mechanical properties of our bilayered neural probes against a hard substrate. Axial compression of beams eventually results in its buckling. The highest force that a sample can withstand before bending is called the buckling force F_buckling_. As depicted in **Fig. 3A**, the compression tests on a hard substrate show clear superiority of the bilayered silk-parylene assembly in terms of mechanical strength. Silk coating improves the probe strength, with an average buckling strength of 10.9 ± 1.3 mN (**Fig. 3D**) (15-times higher than the force required to penetrate the brain-mimicking gel) (**Fig. 2A)**. These experimental values have been compared with predictions of a theoretical analysis (**Fig. 3C**). Our probes were modeled as single beams, whose buckling force is defined by Euler’s formula: F_buckling_ = *π*^*2*^*I*_*x*_*E/(KL)*^*2*^. The buckling force along the x-axis of a clamped beam is linked to its Young’s modulus E, the area moment of inertia I_x_ (*I*_*x*_*= wt*^*3*^*/12*) along the x-axis, L, w, and t are the length, width, and thickness of the probe, respectively and K the column effective length Factor. The cross-section of the device was considered as a constant rectangle and the shanks are beams fixed at one side (K = 0.7). Assuming that E for parylene and silk fibroin are 3 GPa^29, 32^, calculation yields a theoretical buckling force of 7.2 ± 2.3 mN for the probe (**Fig. 3C**). The calculated theoretical buckling force predicts the trend and the order of magnitude of the experimental values (**Fig. 3D**).

**Figure 3.**
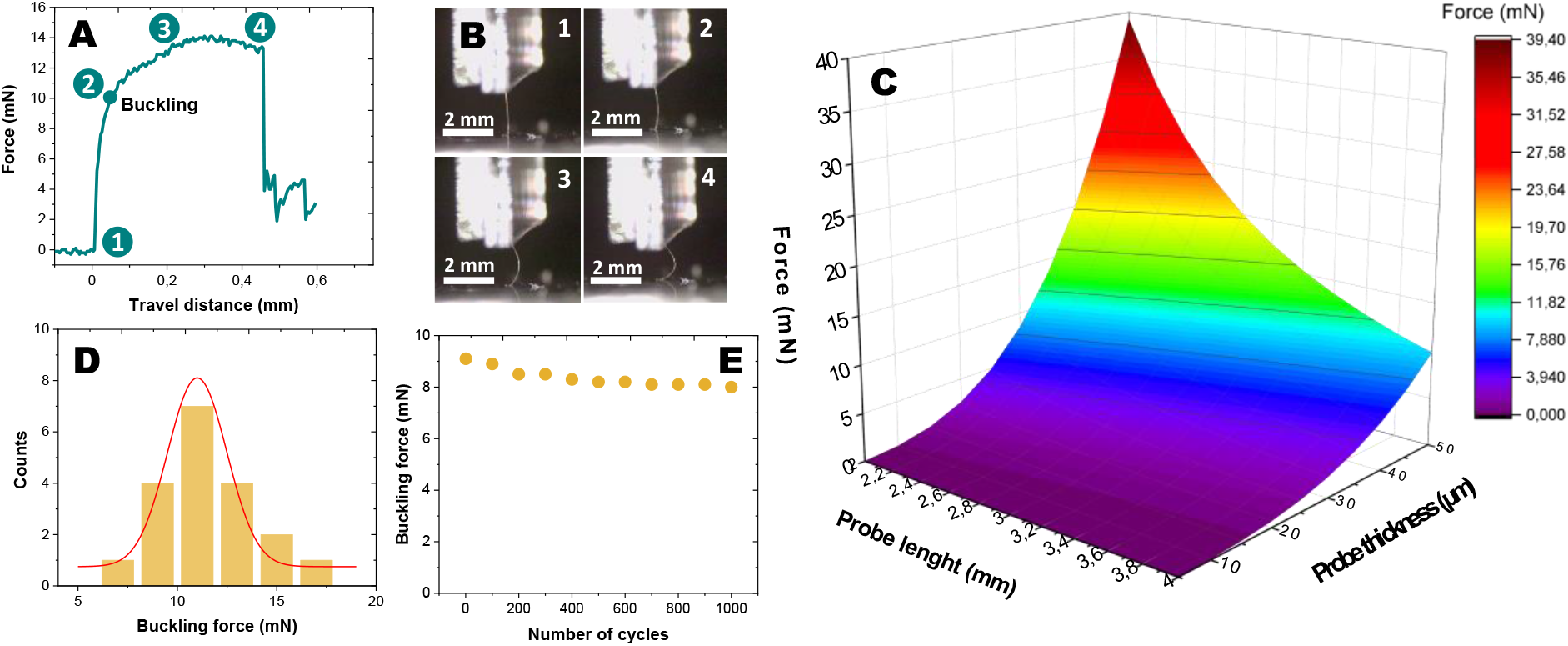
Compression tests of bilayered silk-parylene probe against hard substrate. **A**) Force profile during compression, numbers and buckling position in correspondence with (B). **B**) Series of optical pictures showing the corresponding stages of the shank compression in correlation with the evolution of the force. **C**) Theoretical influence of the shank length (in mm) and thickness (in µm) on the buckling force. The model is a clamped–pinned, the cross-section of the device was considered as a constant rectangle and the shanks are beams fixed at one side and pinned at the other one (K = 0.7). We assumed that E for Parylene C and silk fibroin is 3 GPa. **D**) Electrode-buckling force histogram of the silk-parylene probes (N=20). **E**) Monitoring of the average buckling force as a function of bending cycles with a bending radius of 0.3 mm (1000 cycles).

In addition to axial compression tests, the mechanical stability of our flexible penetrating probes under long-time cyclic compression was measured under cyclic loading with a bending radius of 0.3 mm. As shown in **Fig. 3E**, the original buckling force (first cycle) was measured to be 9.1 ± 0.5 mN, and showed minimal change (only 10% decrease) after nearly 1000 bending cycles. This indicates that such engineered bilayered silk-parylene probes could endure a very large number of contact cycles without any damage.

### Electrical/electrochemical characterizations

Possible alteration of the electrical and/or electrochemical properties of the penetrating probe should also be investigated to ensure their inherent electrode performances. We evaluated the electrical properties of the microelectrodes by electrochemical impedance spectroscopy (EIS), cyclic voltammetry (CV), and calculations of charge storage capacity (CSC). **Fig. 4A** and **B** show the Bode plots across frequencies of interest (10 Hz–7 MHz), for a gold microelectrode of 40 µm in diameter, before and after electrochemical deposition of PEDOT:PSS. The mean impedances at 1 kHz were used for comparison purposes as action potentials have a characteristic frequency band centered at that frequency (**Fig. 4E)**. Before PEDOT: PSS deposition, the average impedance was 210 ± 8.2 kΩ (n = 5) in saline solution, while after polymer deposition, the mean impedance fell to 9.4 ± 0.9 kΩ. This well know-phenomenon is due to a significant increase in the effective surface area with the formation of PEDOT:PSS material, leading to a decrease in impedance of the microelectrode^33^. The corresponding phase plot of the impedance revealed that the PEDOT:PSS-microelectrode was capacitive in the low frequency range (10 Hz) and more resistive at higher frequencies (**Fig. 4B**). The Nyquist plot recorded in saline is presented in **Fig. 4C**. The deposition of PEDOT:PSS produced very small radius of the semi-circle on the Nyquist plot with a charge transfer resistance of about 7.8 kΩ, revealing the low electron-transfer resistance associated with the polymer coating.

**Figure 4.**
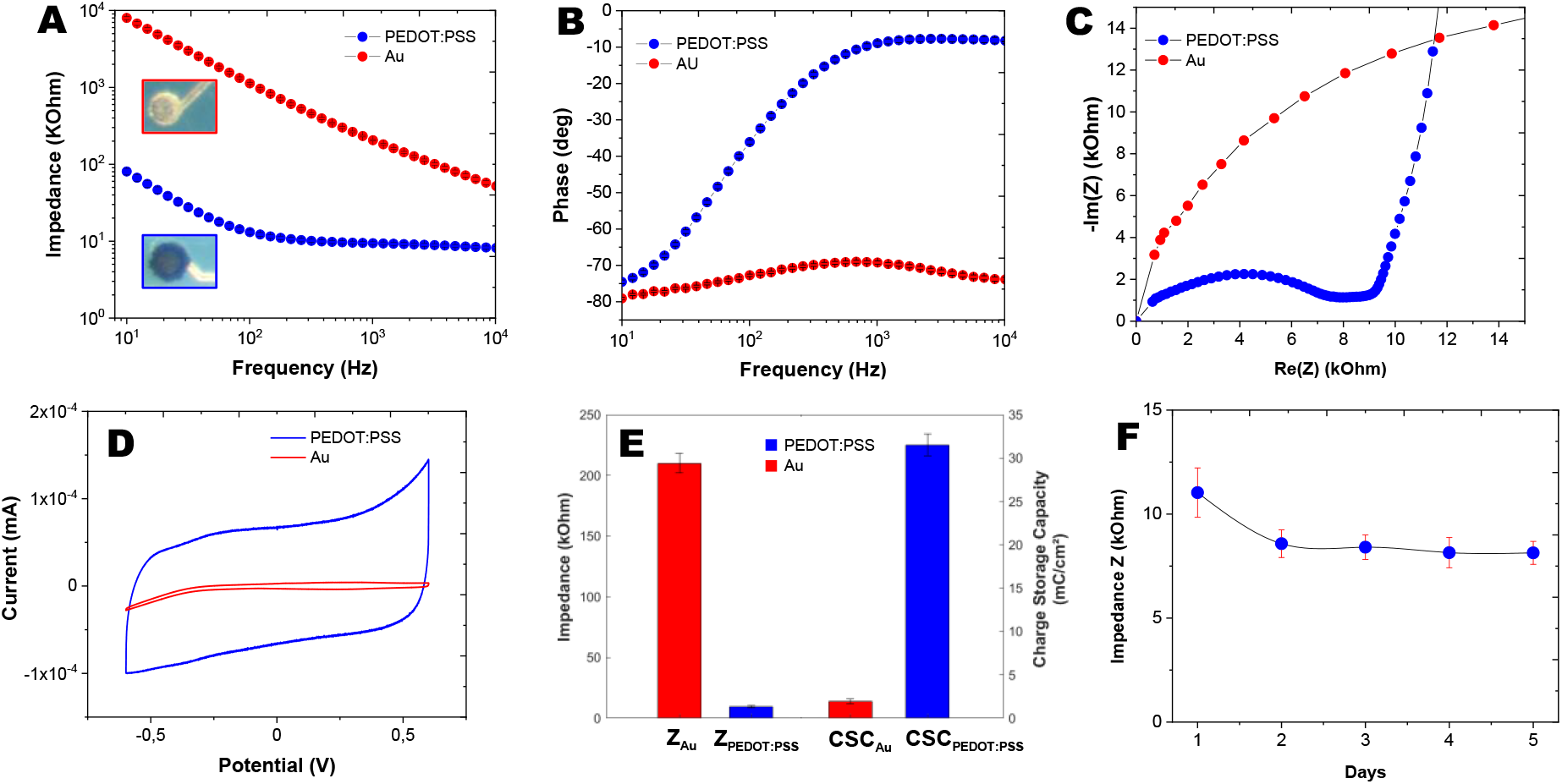
Electrochemical characterizations of microelectrodes. **A**) EIS measurements in PBS buffer at room temperature at frequencies ranging from 10 Hz to 7 MHz of gold and PEDOT:PSS coated microelectrodes. **B**) The corresponding phase vs. frequency plotting of gold and PEDOT:PSS coated microelectrodes. **C**) Nyquist diagram at frequencies ranging from 10 Hz to 7 MHz of gold and PEDOT:PSS coated microelectrodes. **D**) CV in PBS buffer at room temperature by potential sweeping between -0.6 V and +0.6 V *vs*. Ag/AgCl reference electrode at 200 mV/s of gold and PEDOT:PSS coated microelectrodes. **E**) Comparison of the electrochemical characteristics (Z at 1 kHz and CSC) between gold and PEDOT:PSS coated microelectrodes. **F**) Impedance evolution at 1 kHz of PEDOT:PSS coated microelectrode in PBS-soaked gel, performed by EIS measurements.

We next evaluated the charge transfer capabilities of the microelectrodes. A CV with a scan of potential between -0.6 V and 0.6 V at a scan rate of 0.2 V/s was performed and the cathodal CSC was calculated by the time integral of the cathodal currents within the cycled region (**Fig. 4D**). The cathodic CSC increased from an average value of 1.95 ± 0.3 mC.cm^−2^ for the gold to 31.5 ± 1.3 mC.cm^−2^ for the PEDOT-coated microelectrodes. Higher charge capacity may result in higher charge injection that is desirable for neural stimulation.

In addition, to track the integrity of the probe structure and the functional electrical connections, in vitro EIS measurements were carried out daily in saline brain phantoms (gels soaked in PBS buffer). **Fig. 4F** shows the impedance measured at 1 kHz after the insertion of the silk-parylene probe in PBS-based gel as a function of immersion time. The impedance was measured daily from four different probes. Measurements over the 5-day period demonstrated a small decrease followed by a stabilization of the impedances. As the probe was inserted into the gel, we observed no distortion of the probe, which showed that the thickness of the silk layer was sufficient to deliver the full ultrathin structure without compromising the electrical integrity of the parylene probe. The stability in electrical impedance throughout the whole process proves the robustness of our design and protocol for *in vivo* implantation.

### In vitro proteolytic degradation

It is worth mentioning that the bare parylene shank, being only 4 µm thick, curled and could not be handled or manipulated without the bioresorbable silk polymer support. This is shown in **Fig. 5**, where the structural stability of the bilayered probes was analyzed through incubation studies in PBS buffer containing the enzyme protease XIV. The silk biodegradability was measured as the loss of weight of the implant during continuous incubation at 37°C in the proteolytic solution for one week. In a proteolytic environment, the bilayered silk-parylene probes demonstrate a gradual decrease in mass, corresponding to a slow protein fragmentation during incubation. This experiment shows the importance of using the biodegradable silk coating as a temporary stiffener to deliver ultrathin parylene-based flexible devices in deep tissue.

**Figure 5.**
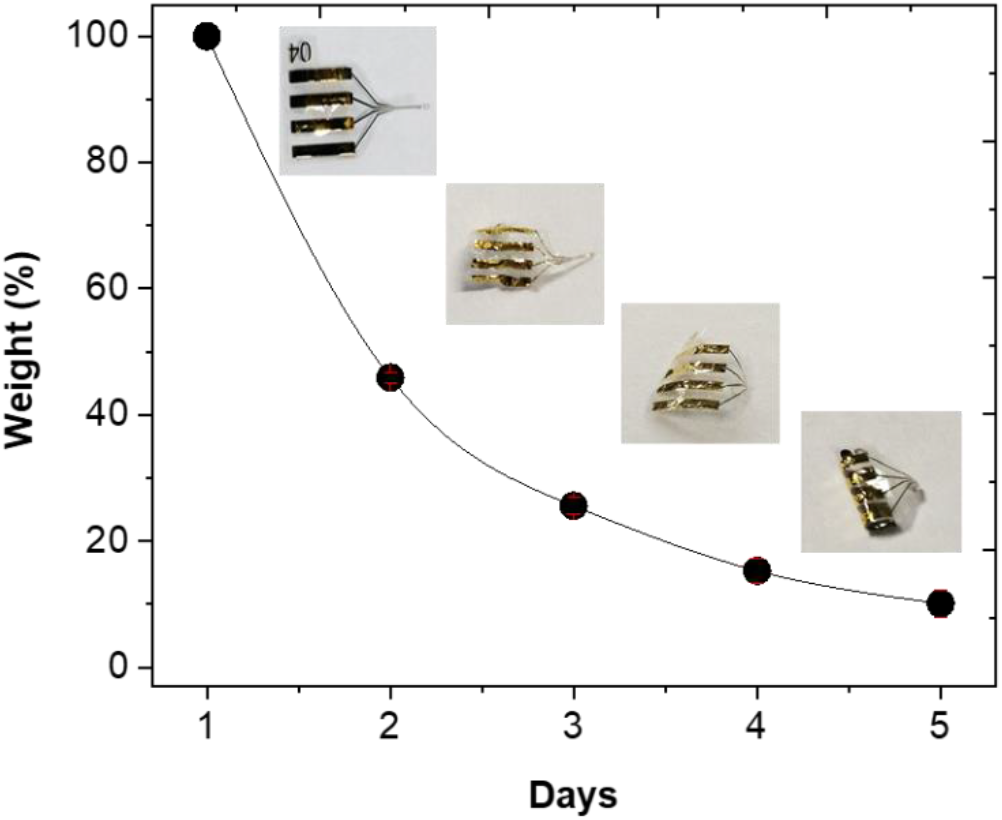
Enzymatic degradation of the bilayered silk-parylene flexible probe in protease XIV (PXIV) solution (1U/ml) during 5 days at 37°C.

### Electrophysiological recording from mouse brain slices

Tests of the probes for their capability to record neuronal activity were made on mouse brain slices. Probe insertion was aimed at layer 2 of the piriform cortex (**Fig. 6A** and **B**). This layer is characterized by a high density of neuronal cell bodies. As spontaneous activity is low or null in most regions of the mouse brain *in vitro*, we added 4-aminopyridine (4AP, 100 µM) to the superfusion solution to activate the slices. 4AP infusion usually leads to the appearance of spiking activity and eventually induces epileptiform activity. The latter provided an opportunity to report on the capability of the probe to record local field potentials in addition to single- and multiunit spiking activity. Recordings followed the implantation of the probe by a few minutes, that is, before the silk coating had enough time to dissolve. These tests were therefore essentially made to ensure that the microelectrodes of the probes allowed for recording of neuronal activity with a good signal-to-noise ratio. Example of spiking activity is presented on **Fig. 6C**.

**Figure 6.**
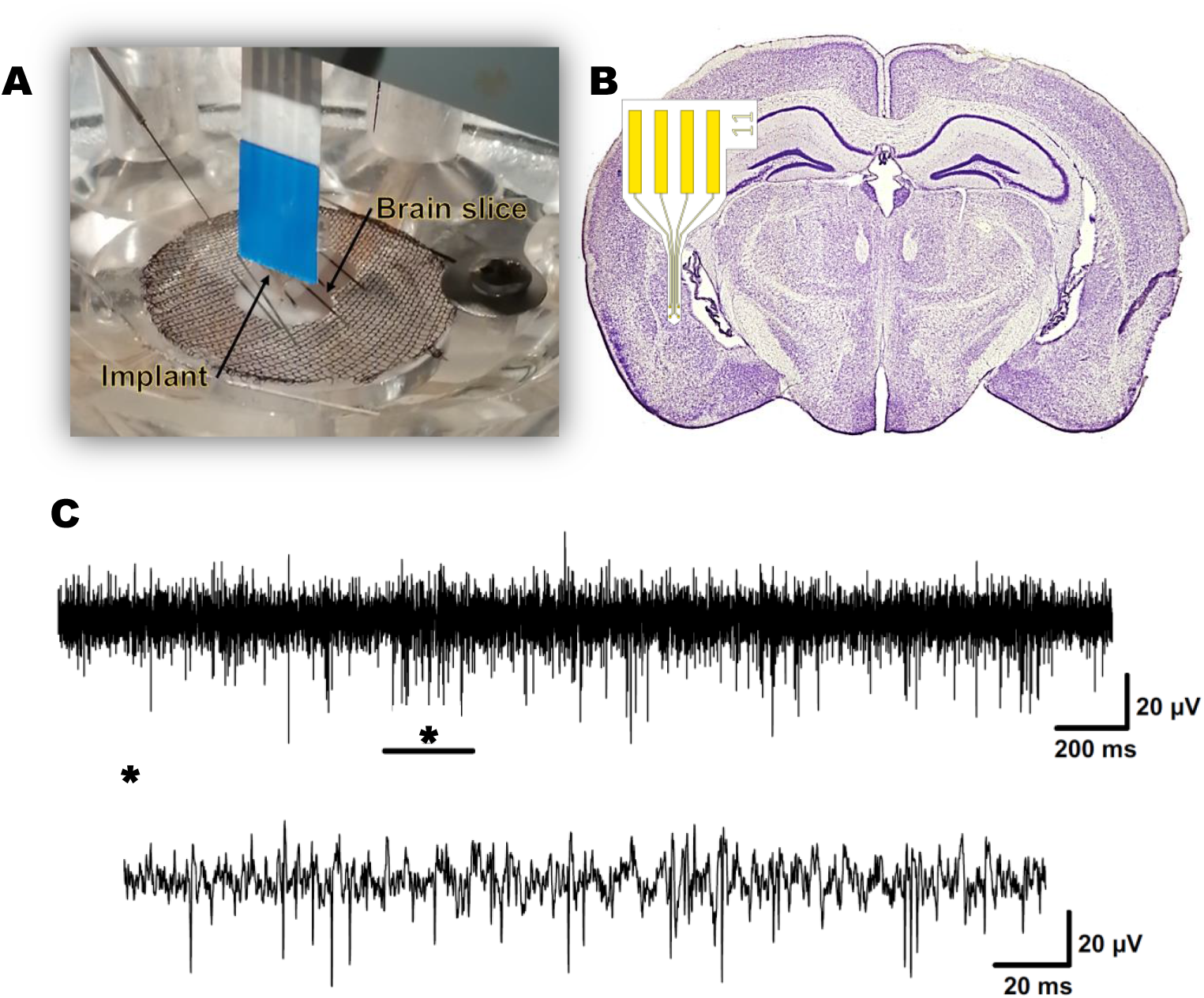
Electrophysiological recording from mouse brain slices. **A**) Photography a 35 μm thick silk parylene probe inserted into a mouse brain slice: insertion occurred with no sign of buckling. **B**) Scheme of the placement of the silk-parylene neural probe. **C**) Electrophysiological signals for PEDOT-coated microelectrodes.

## Conclusions

This work proposes a novel and versatile approach to fabricate, pattern, and deliver ultra-thin penetrating neural probe consisting of a silk-parylene bilayer. The biodegradable silk layer provides a temporary stiffener that can be used to deliver ultrathin parylene-based flexible devices in deep tissue. This innovative and yet robust batch fabrication technology allows complete design freedom of the neural probe in terms of materials, size, shape and thickness. We have systematically studied the behavior of the bilayered structure in brain phantoms and demonstrated that parylene probes as thin as 4 μm can be delivered accurately at desired depth with intact

geometry and electrical functionality. The *in vivo* functionality of our probes remains to be examined in a following study, but our *in vitro* implantation in mouse brain slices for neural recordings suggest that our design provides a novel solution for implanting ultra-flexible and ultrathin devices in the brain while maintaining minimal rejection. In our future works, we envision to correlate the *in vivo* neural recording quality with immunohistochemistry for glial scar formation followed by comparison of the results against existing probes.

## Materials and methods

### Chemicals

Parylene C dimer (PXC) was purchased from Comelec SA. ECI 3012 photoresists was purchased from Microchemicals GmbH. MF CD26 developer was purchased from MicroChem. 3,4-ethylene dioxythiophene (EDOT), Poly (sodium 4-styrenesulfonate) (NaPSS) and acetone were purchased from Sigma Aldrich. Phosphate Buffered Saline (Gibco DPBS 1X) was purchased from Fisher Scientific. Platine (Pt) and Silver chloride (Ag/AgCl) electrodes were purchased from WPI. Solutions were prepared with Deionized water (18 MΩ). Electrochemical characterizations were carried out using a 3-electrode system including a Pt wire as counter electrode, an Ag/AgCl wire as reference electrode and a gold electrode from silk-parylene probes as working electrode.

### Bioresorbable silk fibroin solution preparation

Silk fibroin aqueous solution was prepared from Bombyx mori cocoons following a protocol detailed in SI and previously^30^. The fibroin protein was firstly extracted from the silk fibers by boiling the *silk* cocoons in a solution of 0.02 M Na_2_CO_3_ for 30 min. The regenerated silk fibroin was then recovered and rinsed thoroughly in deionized water before being dried overnight at ambient conditions. The dried silk fibroin was dissolved in a 9.3 M LiBr solution at 60 °C for 4 h. The salt was then removed by dialyzing the solution against deionized water for 24 h at room temperature using a dialysis membrane (MWCO 3.5 KD, Spectra/PorTM) and changing water regularly. The recovered silk fibroin solution had a final concentration of 7% wt.

### Neural probe microfabrication

A 4 inches glass wafer was used to prepare the overall process. The glass substrate was first cleaned by MW-oxygen plasma (800 W, 10 min, O_2_) prior processing. The fabrication began with the deposition of a cellulose acetate layer (≈2 μm) by spin-coating at 1000 rpm for 30 s (5 % w/vol in acetone). It acted as a sacrificial layer to release the final device from the substrate. The silk fibroin aqueous solution (7% wt.) was deposited by drop casting and left drying at ambient conditions overnight, resulting in a 30-µm thick silk film. The thickness of the resulting film is controlled by adjusting the volume of the silk fibroin solution. Then a 3 μm-thick base layer of Parylene C (PXC) was deposited onto the silk-coated substrate through CVD using a C30S Comelec equipment. A 50/200 nm layer of Ti/Au is then deposited by evaporation and patterned through an electroplated Nickel-based shadow mask. Another 1.3 μm top layer of Parylene C was deposited onto the processed metal layer. The shape of the electrode pads, contacts and device body were defined by photolithography steps followed by reactive ion etching in O_2_/CF_4_ plasma (75/25) 500 W and 20 mT. Finally, the dissolution of the cellulose acetate sacrificial layer in acetone was achieved to release the bilayered silk-parylene probes.

### Electrochemical characterizations

To demonstrate the proper electrical functioning of the electrodes, standard electrochemical characterizations were conducted. Electrochemical Impedance Spectroscopy (EIS) and Cyclic Voltammetry (CV) methods were performed using a Bio-Logic VMP3 potentiostat. CV was performed in Phosphate Buffered Saline (Gibco DPBS 1X) at room temperature by potential sweeping between -0.6 V and +0.6 V vs. Ag/AgCl reference at 200 mV/s, allowing cathodal Charge Storage Capacity (CSCc) evaluation^34-35^. EIS was also performed in PBS at room temperature by applying 10 mV sine wave at frequencies ranging from 10 Hz to 7 MHz. The improvement of the electrical properties was achieved by PEDOT:PSS deposition. CV was performed in EDOT: NaPSS solution (10 mM:34 mM) at room temperature by potential sweeping between -0.7 V and +1 V *vs*. Ag/AgCl reference at 10 mV/s. CV and EIS were then performed again to compare the evolution of CSCc, impedance, phase and Nyquist.

### Standard compression tests

Standard compression tests against a hard substrate were performed to assess the mechanical properties of the bilayered silk-parylene probes. The axial compression of the implant allows to evaluate its buckling force, which corresponds to the highest force that a sample can withstand before bending. To guarantee the stability of the test, the implants were held between two glass plates and the compression was carried out over a tip length of approximately 3.4 mm. The silk-parylene probes were fixed to on a MARK-10 ESM303 test bench coupled with a MARK-10 M5-012 force sensor. A Labview program developed internally allows the equipment to be controlled and the force to be monitored according to the displacement of the implant. Compression tests were performed with a slow speed of 2 mm.min^-1^ for optimal monitoring of the buckling force. A video recording of the compression test using a camera Dino-Lite Edge was made to complete the experiment.

### Insertion testing on brain phantoms

Insertion test of the flexible probes was performed using 1% w/v agarose gel brain phantoms imitating the mechanical properties of the brain tissue. In the same way as the compression tests, the silk-parylene probes were held between two glass plates and fixed to the MARK-10 test bench. In order to limit the damage during the insertion tests and to have an optimal monitoring of the forces involved, the experiment was carried out at a very low speed of 0.5 mm.min^-1^. A video recording was made using the DINO-LITE Edge camera to observe the different stages of insertion in correlation with the evolution of the force.

### Dissolution testing

To assess the biodegradation of the silk fibroin layer, an enzymatic degradation test was carried out over several days. 5 implants (∼2 mg) were selected and incubated in 1 ml of protease (Proteas XIV from S.griseus, 3.5 U/mg, Sigma-Aldrich) and PBS solution (1 U/ml of PBS buffer) at 37°C. The implants were photographed and weighed every day after being cleaned with DI water and dried at 60°C for 10 min. The enzyme solution was changed after each weighing to maintain the activity.

### Brain slices preparation

All procedures were conducted in accordance with the guidelines from the French Ministry of Agriculture (décret 87/848) and from the European Community (directive 86/609) and was approved by the ministère de l’Enseignement supérieur, de la Recherche et de l’Innovation (N° 15226-2018052417151228). Two- to 4-month-old C57BL/6 wildtype female mice were used for brain slice preparation. The protocol has been detailed previously^36^. Prominent facilitation at beta and gamma frequency range revealed with physiological calcium concentration in adult mouse piriform cortex in vitro and is briefly summarized here. After deep anesthetization with isoflurane, mice were rapidly decapitated. The brain was removed and prepared for slicing in ice-cold modified ACSF (mACSF). The composition of the mACSF was (in mM): NaCl 124, NaHCO_3_ 26, KCl 3.2, MgSO_4_ 1, NaH_2_PO_4_ 0.5, MgCl_2_ 9, Glucose 10. The mACSF was bubbled with a gas mixture of 95% O_2_ and 5% CO_2_. Four hundred-micrometer-thick slices were cut on a vibratome in the presence of cold, oxygenated mACSF. Slices were allowed to recover for at least one hour at room temperature in a holding chamber filled with oxygenated, *in viv*o-like ACSF, whose composition was (in mM): NaCl 124, NaHCO_3_ 26, KCl 3.2, MgSO_4_ 1, NaH_2_PO_4_ 0.5, CaCl_2_ 1.1, and glucose 10. This ACSF was continuously bubbled with a 95% O_2_ / 5% CO_2_ mixture (pH 7.4).

### Electrophysiological recordings

For recording, a slice was transferred to a submersion type chamber that was continuously gravity-fed with oxygenated (95% O_2_ / 5% CO_2_) *in vivo*-like ACSF at a flow rate of 3-3.5 ml/min. All recordings were performed at 34-35°C. Electrophysiological signals were amplified (×1000) and filtered (0.1 Hz-10 kHz) with a NeuroLog system (Digitimer, UK). Fifty Hz noise was eliminated with a Humbug system (Quest Scientific, Canada). Bandpass filtering (300-3000 Hz) was used to isolate spiking activity online. All signals were digitized with a digitization rate of 50 kHz (power1401, CED, UK). Real-time display of the signals was achieved with an oscilloscope and Spike2 software (CED, UK).

## Acknowledgments

The authors acknowledge fundings from the Agence Nationale de la Recherche (ANR-15-CE19-0006 and ANR-19-CE19-0002-01). This work was supported by French RENATECH network.

## Authors details

^1^LAAS-CNRS, department of Micro Nano Bio Technologies (MNBT), 7 avenue du Colonel Roche, F-31400, Toulouse, France.

^2^CerCo, Université Toulouse 3, CNRS, Pavillon Baudot, CHU Purpan, BP 25202, 31052 Toulouse, France

