## Supplementary Material for "Scalable Batch Fabrication of Ultrathin Flexible Neural Probes using Bioresorbable Silk Layer"

### Bioresorbable silk fibroin solution preparation

Silk fibroin aqueous solution was prepared from Bombyx mori cocoons. The cocoons (5 g) were opened in half and the silkworm was removed before being cut into small pieces (**Fig S1.A**). The sericin protein was firstly removed from the silk fibers by boiling the *silk* cocoons in a 1L beaker containing solution of 0.02 M $Na_2CO_3$ for 30 min. The regenerated silk fibroin was then recovered and rinsed thoroughly in deionized water for 1h before being dried overnight at ambient conditions. The dried silk fibroin (3.6 g) was dissolved in a 50 mL beaker containing 9.3 M LiBr solution at 60 °C for 4 h. The salt was then removed by dialyzing the solution against deionized water for 24 h at room temperature using a dialysis membrane (MWCO 3.5 KD, Spectra/PorTM) and changing water regularly. The recovered silk fibroin solution (**Fig S1.B**) had a final concentration of 7% wt. A centrifugation was performed in order to remove impurities. The solution was placed in a 50 mL tube and centrifuged at 4500 g for 30 minutes at 4°C, the operation was repeated twice by changing the tube between two cleanings.

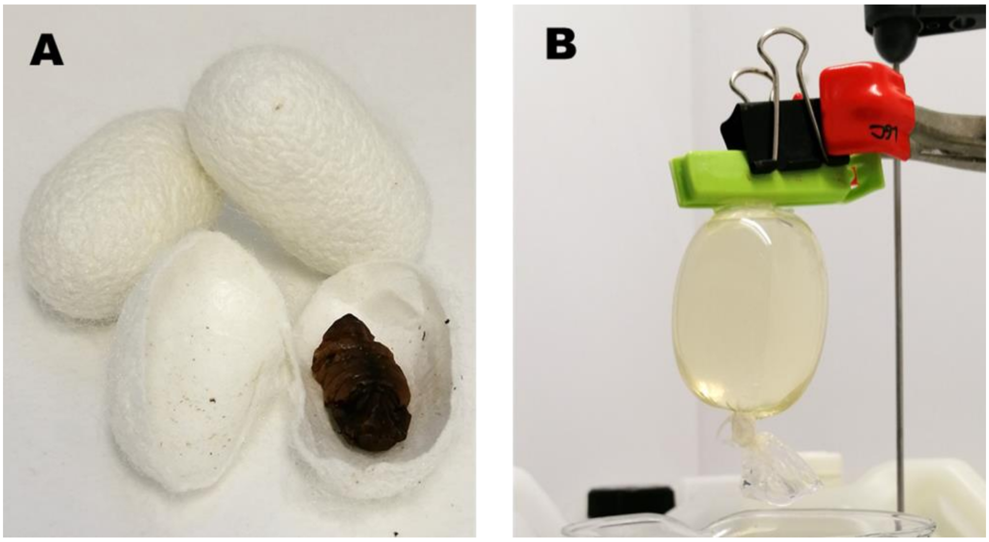


**Figure S1. Preparation of silk cocoons for the production of silk fibroin solution. A)** Opening of silk cocoons with a silkworm inside. **B)** Recovery of silk fibroin solution after 24 hours of dialysis.

### Neural probe microfabrication

A 4 inches glass wafer was used to prepare the overall process to be compatible with standard lithography techniques and clean room procedures. The glass substrate was first cleaned with in a Tepla machine by MW-oxygen plasma (800 W, 10 min, $O_2$) prior processing. The fabrication began with the deposition of a of cellulose acetate layer (≈2 µm) by spin-coating at 1000 rpm for 30 s (5 %wt.vol in acetone solution) (**Fig S2.1**). It acted as a sacrificial layer to release the final device from the substrate. The silk fibroin solution (7% wt.) is deposited by drop casting and left drying at ambient conditions overnight, resulting in a 30-µm thick silk film (**Fig S2.2**). The thickness of the resulting film is controlled by adjusting the volume of the silk fibroin solution. Then a 3 µm-thick base layer of Parylene C (PXC) was deposited onto the silk-coated substrate through CVD using a C30S Comelec equipment (**Fig S2.3**). A nickel shadow mask was placed on the surface of the Parylene (**Fig S2.4 & Fig S3.A**) and held in place with a magnet. A 50/200 nm layer of Ti/Au was then deposited by evaporation (**Fig S2.5**) and patterned through the shadow mask (**Fig S2.6**). Another 1.3 µm top layer of Parylene C was deposited onto the processed metal layer (**Fig S2.7**). The shape of the electrode pads and contacts were defined by photolithography steps using a 2.6 µm ECI resist (**Fig S2.8**) followed by reactive ion etching in $O_2$ plasma 500 W and 20 mT (**Fig S2.9**). The shape of the device body was defined by photolithography step using a 20 µm NLOF resist (**Fig S2.9**) followed by reactive ion etching in $O_2/CF_4$ plasma (75/25) 500 W and 20 mT. Finally, the dissolution of the cellulose acetate sacrificial layer in acetone was achieved to release the bilayered silk-parylene probes (**Fig S2.11 & Fig S3.B**).

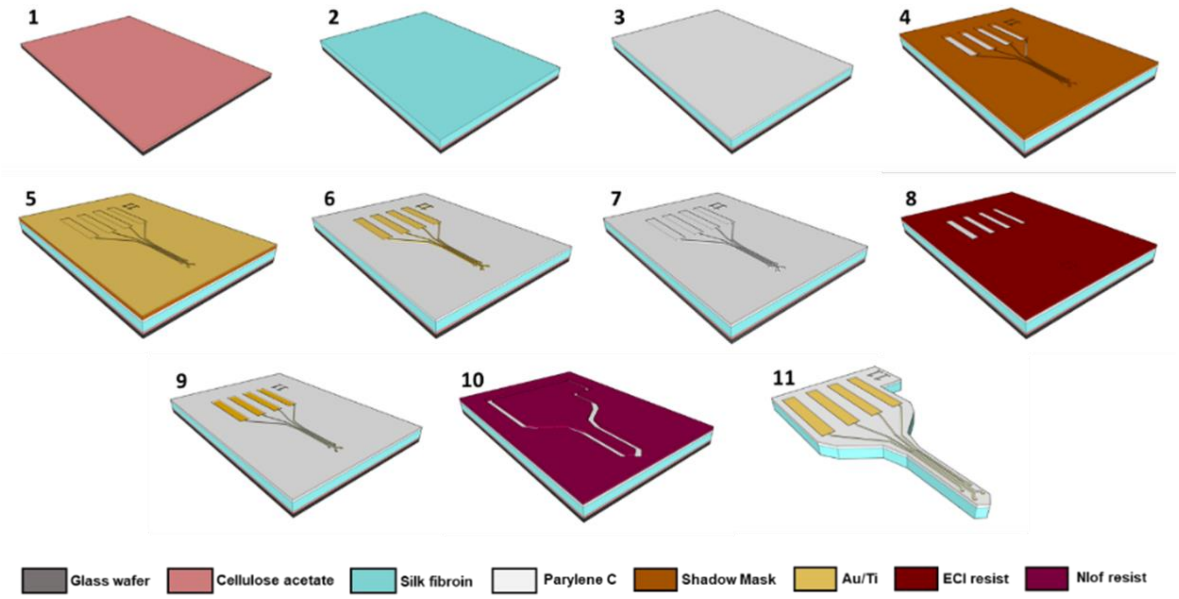


**Figure S2.** Schematic illustration of bilayered silk-parylene neural probes fabrication.

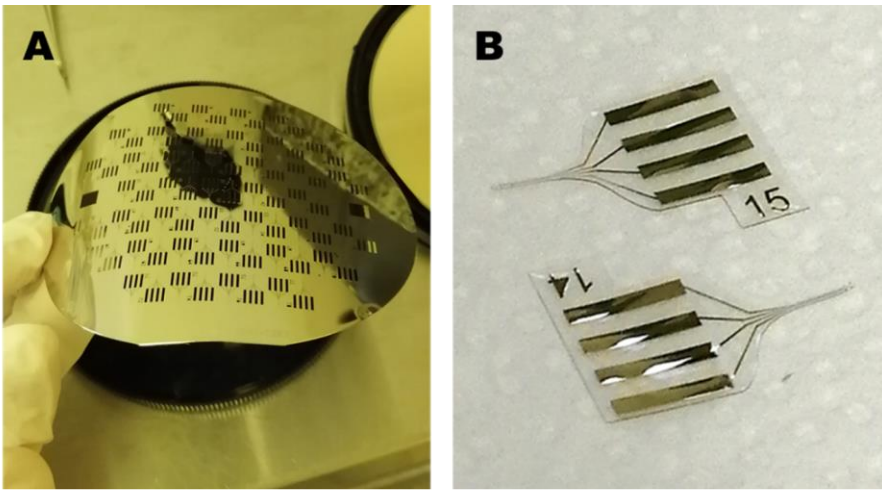


**Figure S3. Silk-parylene neural probes fabrication overview. A)** Picture of the shadow mask used for metallization. **B)** Picture of two bilayered silk-parylene neural probes**.**

### Electrochemical cleaning and deposition

To demonstrate the proper functioning of the electrodes and to characterize them, an electrochemical study was conducted. Before utilization, the electrodes must be electrochemically cleaned to ensure good deposition conditions. First, a series of 20 potential pulses (2 V vs. Ag/AgCl for 1s, -1 V vs. Ag/AgCl for 1s) was applied in PBS at room temperature. To probe the microelectrodes, a CV was performed in PBS at room temperature by potential sweeping between -0.3V and +1.3 V vs. Ag/AgCl reference at 200 mV/s.The activation of the microelectrodes made it possible to find a good electrochemical signature of gold (**Fig S4.A**). The improvement of the electrical properties was achieved by PEDOT:PSS deposition. CV was performed in EDOT:PSS solution (10 mM:34 mM) at room temperature by potential sweeping between -0.7 V and +1 V vs. Ag/AgCl reference at 10 mV/s (**Fig S4.B**).

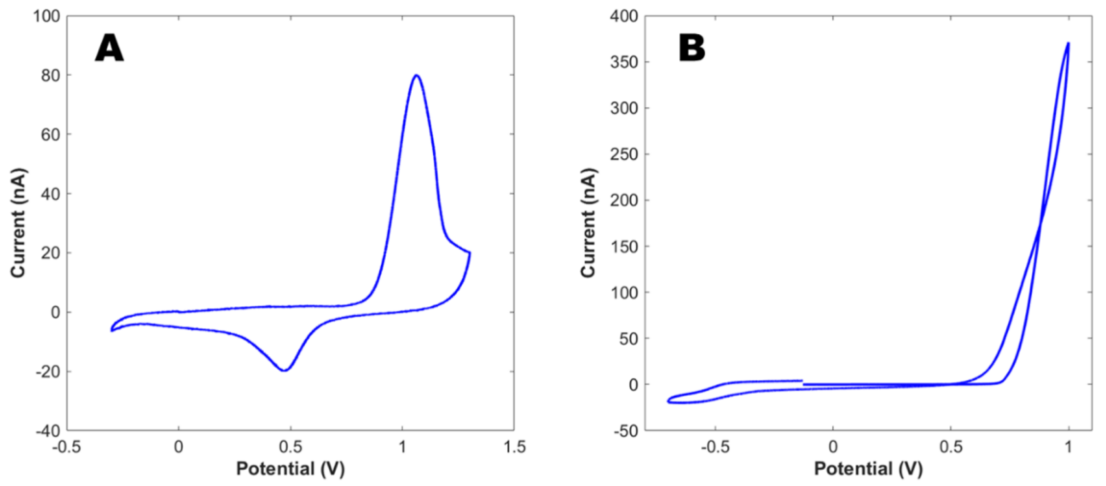


**Figure S4. Electrochemical cleaning and characterizations of neural microelectrodes. A)** CV in PBS at 200 mV/s between -0.3 V and +1.3 V vs. Ag/AgCl reference after activation. B) PEDOT:PSS deposition by CV in EDOT:PSS solution at 10 mV/s between -0.7 V and +1 V vs. Ag/AgCl reference.
